# Female Moths Incorporate Plant Acoustic Emissions into Their Oviposition Decision-Making Process

**DOI:** 10.1101/2024.11.06.622209

**Authors:** Rya Seltzer, Guy Zer Eshel, Omer Yinon, Ahmed Afani, Ofri Eitan, Sabina Matveev, Galina Levedev, Michael Davidovitz, Tal Ben Tov, Gayl Sharabi, Yuval Shapira, Neta Shvil, Maya Harari Gibli, Ireen Atallah, Sahar Hadad, Dana Ment, Lilach Hadany, Yossi Yovel

**Affiliations:** School of Zoology, George S. Wise Faculty of Life Sciences, Tel Aviv University, Ramat Aviv, 6997801, Israel; Plant Protection Institute, Agricultural Research Organization -Volcani Institute, Derech Maccabim 68, Rishon LeZion, Israel; School of Plant Sciences and Food Security, George S. Wise Faculty of Life Sciences, Tel Aviv University, Ramat Aviv, 6997801, Israel; Sagol School of Neuroscience, Tel Aviv University, Ramat Aviv, 6997801, Israel

**Keywords:** plant-insect interactions, ultrasound-hearing moths, plant ultrasonic clicks

## Abstract

Insects rely on plants’ visual, chemical, tactile, and electrical cues when making various decisions. A recent study demonstrated that dehydrated plants emit ultrasonic sounds within the auditory sensitivity range of many moth species. In this study, we sought to determine whether insects also rely on plant acoustic signals when making decisions. We investigated whether female moths rely on ultrasonic clicks which are typically produced by dehydrated plants when deciding where to oviposit. In the absence of an actual plant, the moths indeed preferred to lay their eggs in proximity to acoustic signals which represent dehydrating plants. Tracking the moths’ behavior prior to the decision showed that they examined both sides of the arena and gradually spent more time on the acoustic-playback side. Interestingly, when actual plants were added to the arena, the oviposition preference was reversed and the moths preferred silent plants, which is in accordance with their a-priori preference for hydrated plants. Deafening the moths eliminated their preference, confirming that the choice was based on hearing. Moreover, the presence of male moths including their auditory signals did not affect their oviposition decision, suggesting that the response was specific to plant sound emissions. We reveal evidence for a first acoustic interaction between moths and plants, but as plants emit various sounds, our findings hint to the existence of more currently unknown insect-plant acoustic interactions.

## Introduction

Plant-insect communication has been shown to rely on various modalities, including vision, olfaction, and mechanoreception (Boppré 1978; Kevan and Lane 1985; Gori 1989; Ne’eman 1995; Schiestl 2010; Brito *et al*. 2015; van Dam and Bouwmeester 2016). Plant-insect (airborne) acoustic communication, however, has never been demonstrated. It has long been known that plants vibrate at ultrasonic frequencies due to physiological processes such as cavitation, resulting from changes in their water pressure (Milburn and Johnson 1966; Tyree and Dixon 1983; Ponomarenko *et al*. 2014). Recently it has also been shown that these ultrasonic sounds produced by a drought-stressed or cut plant are airborne and are probably loud enough to be detected by ultrasound-hearing moths from a distance of a few meters (Khait *et al*. 2023). Moreover, it was shown that these sounds can serve as reliable cues for the condition of the plant, specifically indicating whether a plant is drought-stressed.

Ultrasonic hearing abilities and hearing organs located on different body parts have evolved multiple times independently in several Lepidoptera families. Hearing sensitivity typically falls within the 20 kHz −60 kHz range in all groups of moths that have evolved ultrasonic hearing (Fenton and Fullard 1979; Hoy 1996; Conner 1999; Robert and Göpfert 2002; Moir *et al*. 2013; Göpfert and Hennig 2016). Two main hypotheses exist regarding the evolution of these hearing organs. The first suggests that they have evolved for sexual communication, i.e., to detect ultrasonic signals produced by male moths (Nakano *et al*. 2009). The second hypothesis suggests that they have evolved as an anti-predator mechanism to detect echolocation calls produced by bats (Conner 1999; Greenfield and Weber 2000; Nakano *et al*. 2014; but see Kawahara *et al*. 2019). Regardless of why it has evolved, ultrasonic hearing allows moths to detect various additional sounds (Spangler 1988), including plant dehydration sound clicks which have a wide spectrum that overlaps with moths’ hearing range and peaks around 50kHz (Khait et al. 2023). We thus hypothesized that herbivor female moths with ultrasonic hearing might exploit ultrasonic plant emissions as cues to infer plant condition and employ this information for oviposition.

The selection of an oviposition site has a significant impact on the fitness of the hatching herbivor larvae and is thus one of the most critical decisions in the life of a female moth (Lhomme *et al*. 2018). In this study, we examined the Egyptian cotton leafworm (*Spodoptera littoralis*; Noctuidae)—a polyphagous herbivore and one of the most significant pests of tomato plants (Prasad and Bhattacharya 1975), which possesses tympanic ears tuned to ultrasonic frequencies (Tougaard 1996, Skals *et al*. 2005, Anto *et al*. 2011). The ears’ sensitivity of many moths from the Noctuidae family have been fully characterized and they typically show a wide range of sensitivy between ∼20 - ∼60 kHz (Fullard 1998). The full audiogram of the Egyptian cotton leafworm moth has not been documented, but (in accordance with the moths in the Noctuidae family) its hearing has been shown to be most sensitive around 38 kHz, a frequency which is part of the plant’s click spectrum (Tougaard 1998). Moreover, the spectra of the clicks of the males of this species (Fig.1), which are clearly heard by the females broadly overlap with plant clicks. We further demonstrated that the moth can hear echolocation calls which are in the range between 40-80kHz, thus demonstrating sensitivity in the plant clicking range (see Methods).

**Fig. 1:**
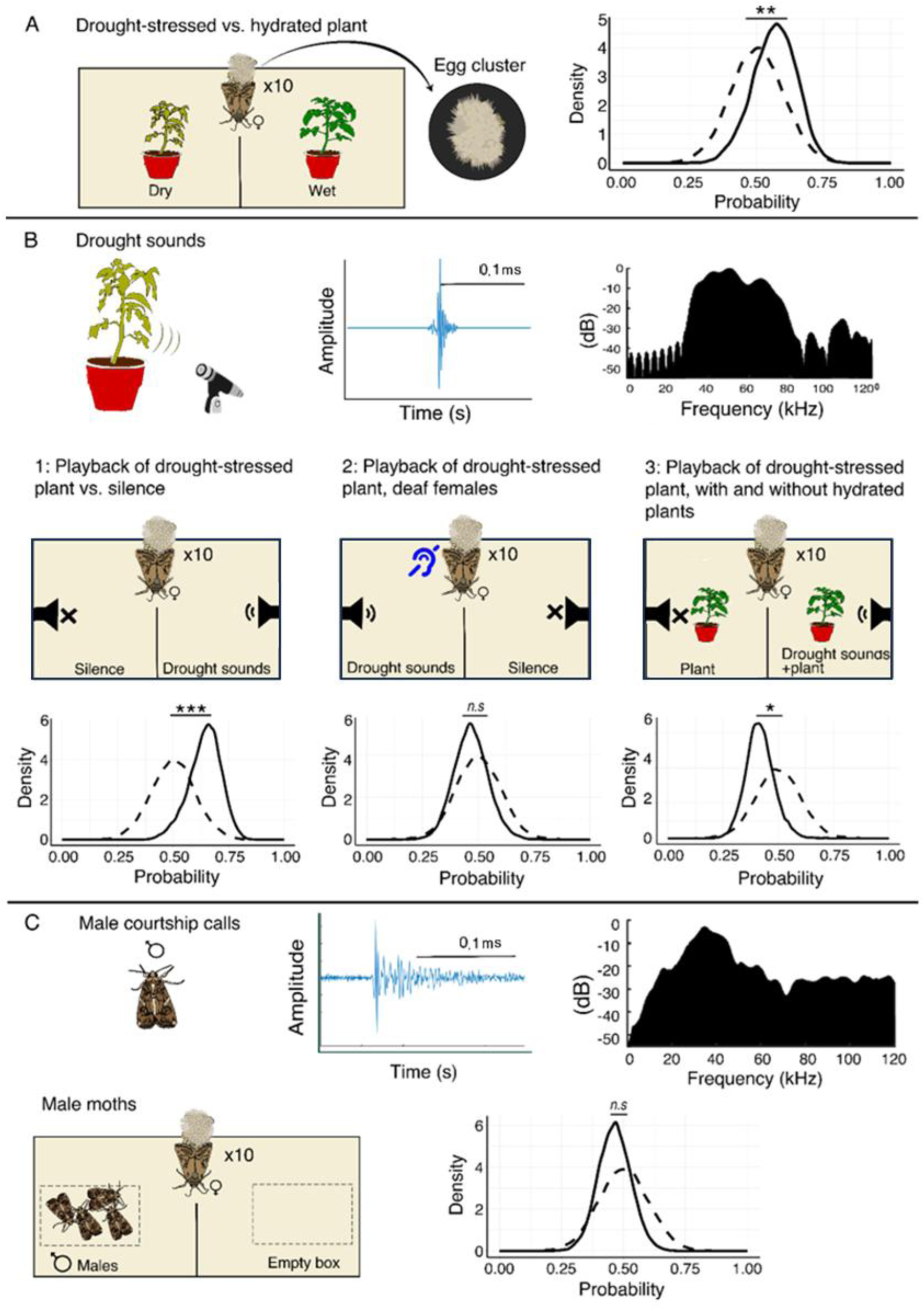
The setup and results. In all panels A-D, the sound played in the setup is presented in the left section (treatment). Because the number of egg clusters were low (between 0-5 clusters) we find that presenting the Bayesian posterior (see Methods) for the probability to lay a cluster is more informative (We present the raw data on Supplementary Figure 2). The posterior distribution is depicted by solid lines. The prior distribution (with a mean of 0.5 and an STD of 0.1) is represented by dashed lines. To create these plots, eggs laid on the tested side (where the speaker was active, or hydrated plant in the initial experiment) are denoted as 1, while those on the opposite side are marked as 0. These plots thus demonstrate the probability of obtaining a 1 or 0 in each experiment. The middle section shows the two-choice oviposition setup, and the right side shows the results for the following conditions: A) Drought-stressed vs. thriving plant (no playback). B1) Silence vs. drought-stressed plant playback (without a plant). B2) Deaf females in a setup with silence vs. drought-stressed plant playback (without a plant). B3) Silent plant vs. playback of drought-stressed plant. C) A box with male moths vs an empty box. Tomato and male clicks are presented (time signal and spectrum) in panels B and C. The horizontal black bar depicts 0.1ms.

Much research has been conducted to characterize the females’ oviposition choice in this species with many factors suggested to be important for their decision-making process. The females have been found to prefer certain species of host plants over others (Salama *et al*. 1971; Sadek *et al*. 2010), to select plants based on their larval experience (Proffit *et al*. 2015), and to choose plants devoid of parasitic larvae, possibly because the presence of such larvae could promote the recruitment of natural enemies (Sadek et al. 2010). Studies have also investigated female preferences in response to plant stress signals, particularly olfactory cues. However, there is no clear consensus on the direction of these preferences (e.g., Chen *et al*. (2008) and Showler & Moran (2003)). Nonetheless, it is widely accepted that females are capable of recognizing and responding to these signals.

In this study, we investigated whether ultrasonic sounds typical of drought-stressed plants influence oviposition decision making in the Egyptian cotton leafworm moths. Based on their general behavioral preference for non-dry plants (as we validated, see below), we hypothesized that the female moths would be affected by plant ultrasonic signals when making oviposition decisions. Our results support this hypothesis, providing the first evidence for the use of typical plant sounds by insects.

## Results

In each of the following experiments, we placed 10.9±0.17 (Mean±SE) fertile female *S. littoralis* moths in the center of a 100×50×50 cm^3^ arena divided in the middle, with two choices offered, one on either side of the arena (a two-alternative forced choice paradigm, see Methods). To assess their choice, we compared the number of egg clusters which the moths had laid on each side.

Each treatment was repeated at least 9 times (i.e., with a new set of moths) but the moths in each repetition were observed for several consecutive nights so that the minimum number of egg-laying events per treatment was 17. Each night was considered an independent observation because the moth could make a new decision regarding where to lay her eggs (to account for this repetition, the nights were nested in the statistical model). The treatment and the control sides were alternated between repetitions. To ensure replicability, the main plant-acoustic treatments were run twice with a pause of several months in between (see Table 1 in the Methods). In these experiments, we used the number of egg clusters, rather than the total number of eggs, as the response variable because each cluster represents a distinct oviposition decision. However, we describe a third experiment below where we evaluated the effect of the plant sounds on egg number (and not cluster number).

**Table 1.** Summary of experimental conditions including the number of repetitions, i.e. the number of times that new moths were placed in the arenas and the number of observations (each repetition was observed for approximately three consecutive nights). The total number of egg clusters and the P-values for each experiment are reported. Experiments that were replicated twice appear in two separate lines denoted for combined statistics and by #1 or #2. Experiments and observations that did not produce any egg-laying were excluded from the data set and that is why the number of observations is often the same as the number of repetitions.

| Experiment | #Repetitions | #Observations | #Egg clusters | Mean± SE clusters on the side of the treatment | Mean ± SE clusters on the side of the Control | P-Value | Estimates (# of Egg clusters) |
| --- | --- | --- | --- | --- | --- | --- | --- |
| Drought-stressed plants vs. well-hydrated plants | 17 | 17 | 53 | 0.88±1.11 | 2.23±2.68 | 0.01 | 0.93 |
| Playback of a drought-stressed plant vs silence, combined trials (playback: 60 per minute) | 38 | 45 | 67 | 1.08±0.82 | 0.40±0.65 | 0.00 | 1.00 |
| Playback of a drought-stressed plant vs silence 1# (playback: 60 per minute) | 11 | 17 | 24 | 1.11±0.69 | 0.29±0.58 | 0.01 | 1.34 |
| Playback of a drought-stressed plant vs silence2# (playback: 60 per minute) | 27 | 28 | 43 | 1.07±0.89 | 0.46±0.69 | 0.02 | 0.84 |
| Deafened moths -Playback of a drought-stressed plant vs silence | 23 | 23 | 39 | 0.70 ± 0.70 | 1.00 ± 1.09 | 0.55 | 0.12 |
| Well-hydrated plants and playback of a drought-stressed plant, combined trials (playback: 60 per minute) | 29 | 39 | 110 | 1.05±0.99 | 1.76±1.64 | 0.01 | -0.52 |
| Well-hydrated plants and playback of a drought-stressed plant 1# (playback: 60 per minute) | 9 | 19 | 44 | 0.78±0.91 | 1.52±1.38 | 0.05 | -0.66 |
| Well-hydrated plants and playback of a drought-stressed plant 2# (playback: 60 per minute) | 20 | 20 | 66 | 1.30±1.03 | 2.00±1.86 | 0.10 | -0.43 |
| Males vs no-males | 19 | 29 | 48 | 0.72±0.92 | 0.93±1.33 | 0.39 | -0.25 |

First, to examine whether *S. littoralis* females prefer to lay their eggs on drying or fresh tomato plants (without any playback sound, see Exp. 1 in the Methods), we placed them in an arena with one drying and one fresh plant. Female *S. littoralis* demonstrated a strong preference to lay their eggs on fresh plants that were not drought-stressed (Fig. 1A, 2.2±2.7 vs. 0.9±1.1 egg clusters; Mean ± SE; clusters per night respectively, *p* =0.004, Mixed effect generalized linear models – GLMM with the number of egg clusters as the explained parameter, the treatment as a fixed effect and the number of the arena and the repetition round and night as random effects, see statistics).

We next examined whether an ultrasonic acoustic stimulus affects moths’ oviposition decision making. To this end, we played drought-stressed sounds (recorded from a real drying tomato plant) on one side of the arena and either placed nothing on the other side or placed a decoy silent resistor to control for electric field sensing (see Exp. 2 in the Methods). Because we aimed to examine the effect of sound only (without other sensory cues such as visual or olfactory), in this condition, there was no plant in the arena, and we placed a small mesh box wrapped with a paper towel in the center of each side to encourage oviposition (the speaker was under the mesh so that the moth could not sense the vibration directly, only through airborne sounds waves). Female moths significantly preferred to lay their eggs on the side of the arena in which drying plant sounds were played (contradicting the initial observation that they prefer hydrated plants Fig. 1A).

Notably, this experiment was repeated twice - six months apart - and the preference was significant both times (Fig.1B1 1.1±0.8 vs 0.4±0.7 egg clusters per night for the playback and the silent side respectively, Mean±SE *p*=0.0004, estimate=1, GLMM as above, see Table 1 for the results of each session). The average number of egg clusters (1.1 clusters per-night) in this condition was lower than in the baseline condition with a plant (2.2 clusters), but this is reasonable when taking into account that there was no plant in the arena. The playback rate was high with 60 drought clicks played per minute. This is higher than the rate reported for a single young plant, but it is feasible when considering a patch of adult plants as we have demonstrated experimentally (see Methods). Moreover, we repeated this experiment in an improved experimental setup with a lower playback rate of 30 per minute and got the same result – see below.

To make sure that the acoustic signals were the sole influential factor in the moths’ decision-making process, we deafened mated female moths (by puncturing the tympanic membrane located at the thoraco-abdominal juncture using an entomological needle #2, see Methods section) and repeated the experiment (drought-stressed sounds-no plant in the arena). We placed 9.3±1.8 female moths in an arena and monitored their choice of oviposition sites. In accordance with the acoustic hypothesis, the deafened moths did not show any preference in egg laying (Fig.1B2, 0.70 ± 0.70 vs. 1.0 ± 1.09 egg clusters per night, *p* = 0.55, estimate = 0.12, GLMM).

To examine the importance of sound in oviposition decision making under pseudo natural conditions, we placed two hydrated tomato plants - one on either side of the arena - and added a speaker playing back drought emissions sounds on one side and on the other side either a resistor (with the same impedance as the speaker) to control for potential effects of the electric field, or nothing. Interestingly, females showed a significant preference for the silent plant. In this case, the female preference was similar to the initial experiment (without playback) in which the females preferred hydrated plants. The females laid 1.8±1.6 vs. 1.1±1.0 egg clusters per night on the silent and playback sides, respectively. This treatment was also repeated twice over a 12-month period (Fig. 1B3, estimate = −0.52, *p*=0.01, GLMM as above, see Table 1 for the results of each repetition, note that the second repeat was only marginally significant).

To assess whether the moths’ response was specific to plant sounds, we conducted an additional test using male moths that were placed on one side of the arena (in a mesh-box so females could not interact with). The male moths produced courtship clicks with a similar spectral range like tomato clicks (as we validated, Methods). Females showed no significant preference to lay their eggs near male moths (see Supplementary Fig. 1, Fig. 1C, p = 0.4, estimate = −0.25, GLMM as above).

To gain further insight into the moths’ decision-making process, we repeated experiment 2 (Fig. 1B) where drought-stressed sounds were played on one side of the arena without a plant in three additional repetitions (with a total of N=13 females) while videoing and tracking the entire behavior. In these repetitions, eggs were laid only on the playback side of the arena. The continuous tracking showed that most moths (8 of the 13) visited both sides of the arena, crossing sides 4.2 ±5.7 times (Mean±SD) on average during the night (Fig.3 A). Moreover, over time, there was a significant increase in the female moths’ tendency to spend more time in the playback side (Logistic GLMM, *p*<0.004, Fig.3 B.).

The sound gradient experiment: To control for a few of the experimental parameters from the setup shown in Fig. 1, we conducted another experiment testing the main effect of plant sounds on oviposition. This experiment replicated the oviposition site preference between the plant stress sound side and the quiet side, but within a different experimental setup (see Sound gradient experiment in the Methods). Namely, in this experiment we tested a single moth each time, with a lower biological-feasible plant click rate (30 click per minute, for experiment regarding natural click rate see Methods) within a long arena – creating a sound gradient. To this end, we placed a single female moth in a 150 cm long arena. On one side of the arena (location −75, Fig. 2A), a speaker played sounds recorded from a drought-stressed tomato plant (at a rate of 30 clicks per minute). On the other side of the arena (location +75, Fig. 2A), there was a silent resistor. A feeder with 60% sugar solution was positioned at the center (location 0, Fig. 2A). For each egg cluster, we then measured the distance from the center where it was laid and the number of eggs it contained. The results, for both egg and cluster numbers, revealed a clear bimodal distribution with peaks near the feeder and the speaker but not at the silent edge of the arena. Hence, most clusters were laid very close to the feeder or the speaker while no eggs were laid near the resistor (the closest egg was 21cm away, Fig. 2B, C, both egg and cluster number distributions were significantly different from the expected H0 distribution which was estimated using permutation, K-S test, p = 2.2 × 10⁻¹⁶ for the clusters, p = 3.9 × 10⁻^14^ for the eggs, and see Methods and Supplementary Fig. 3). To exclude any potential effect of temporal correlations on egg laying, we have also rerun the statistics when only taking the first night, when the females laid clusters to avoid the desensitization or dependency. This test revealed similar results (D = 0.55, p-= 2.2× 10⁻16). This was thus a third independent validation that females prefer to lay eggs near plant playback and that this behavior is seen both when quantifying the individual egg or the cluster level.

**Fig. 2:**
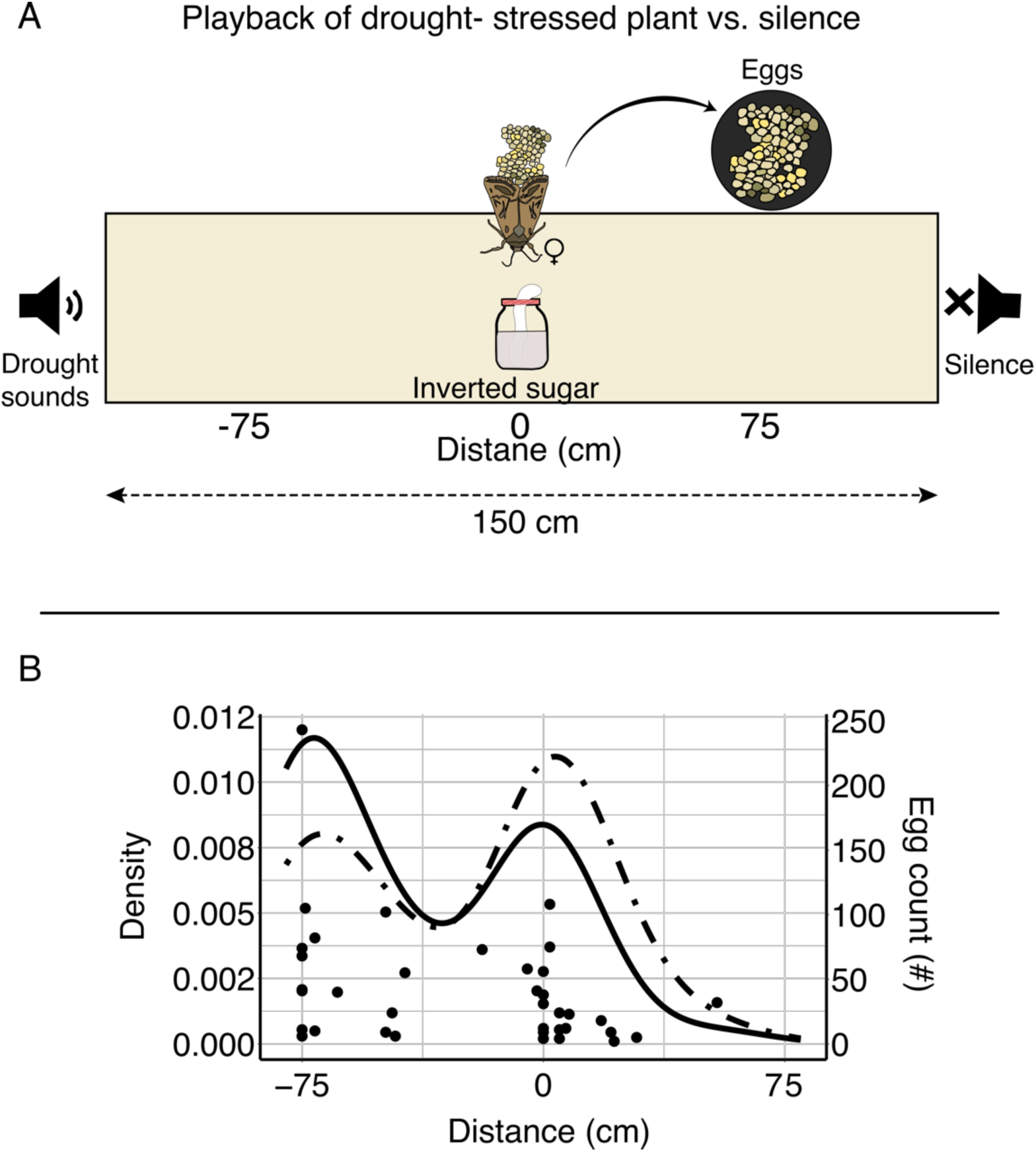
Females lay eggs near acoustic playback. A) The long arena creates an acoustic gradient, allowing us to investigate whether female moths prefer to lay their eggs in specific locations based on the sound environment. Additionally, there is sugar water in the center of the arena, which serves as the adult moth’s food. B) Egg count density (solid line) and cluster density (dashed line). Both figures display a bimodal distribution, with one peak near the speaker (−75) and another near the feeder (0). The points under the graph depict laid clusters, illustrating the relationship between the number of eggs per cluster and their spatial distribution within the arena.

**Fig. 3:**
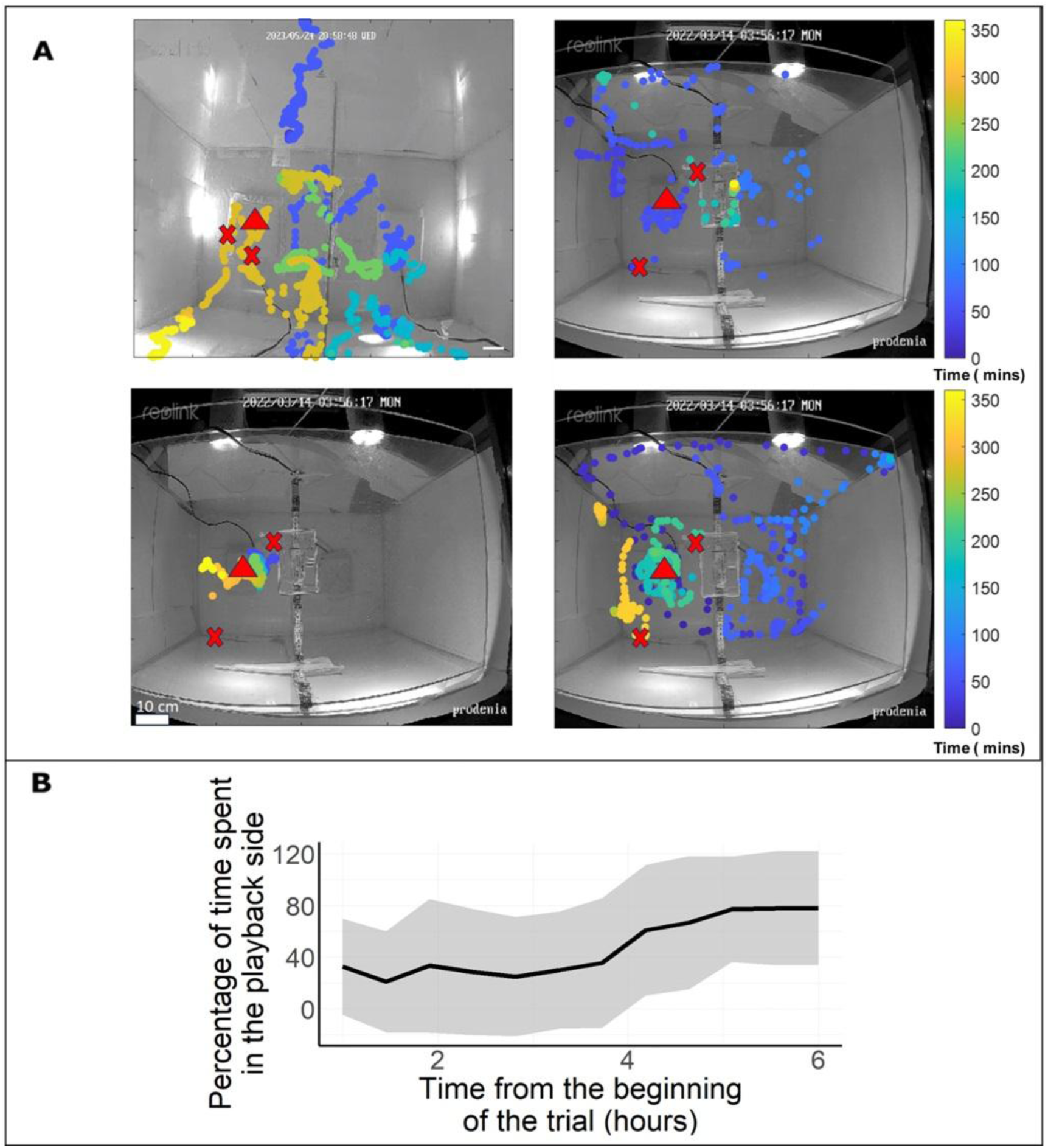
Females’ movement and decision making. A) The continuous location over time in the arena (top-view) of 4 individual moths during one trial of the drought sounds vs. silent treatment. Time is represented by color in minutes, with a red triangle indicating the playback side and red X’s marking the locations where eggs were laid. Note that we cannot be sure which of the individuals laid the eggs. B) The proportion of time moths spent in the playback side (in bins of 30 minutes) increased over time.

As noted, moths prefer to oviposit near stress sounds in a plant-free system (Fig. 1B1), but their response reverses when stress sounds are played in a system containing plants, leading them to choose oviposition sites near the quiet plant (Fig. 1B3). In this experimental system, we aimed to test whether this reversal reflects a general preference for plants (even when stressed) over no-plant options. We offered moths a dehydrating plant with added clicking sounds on one edge of the arena and plain soil on the other (Supplementary Figure 4). Moths significantly preferred to lay their eggs on the dehydrating clicking plant compared to plain soil (Supplementary Figure 4). This experiment conceptually simulates one step prior to Fig. 1B1 - removing the stressed plant while retaining only acoustic signals - suggesting that clicking sounds might be perceived as indicative of plant presence in the absence of multimodal signals.

## Discussion

We reveal first evidence for the use of acoustic information and specifically of sounds typically emitted by plants in insect decision making. Despite decades of research on plant vibrations, it has only recently been shown that these vibrations can be detected remotely by organisms with ultrasonic hearing ability (Khait et al. 2023). Our current results suggest that *Spodoptera littoralis* females detect and respond to ultrasonic clicks which are typically emitted by drought-stressed tomato plants and adjust their choice of oviposition accordingly. This finding opens a whole new range of possibilities for animal-plant acoustic interactions.

Moreover, the presence of clicking male moths had no significant effect on the females’ oviposition preference, suggesting that female moths can distinguish between different sounds and specifically respond to plant-like sounds. Although the moth’s hearing system might be too simple to distinguish among the spectral properties of the different sounds, i.e., male clicks vs. plant sounds (Nakano *et al*. 2013), the temporal patterns of the sequences emitted of these sources are very different. While male moths emit bursts of several clicks (Supplementary. Fig.2), plants emit sporadic clicks with no clear temporal order (as used in our playback). Playback of additional sound signals are needed to examine moth specificity.

Although females responded in both treatments when ultrasonic drought-stressed signals were played, they exhibited opposite preferences depending on the presence of a plant. When there was no plant in the arena, the moths showed a strong preference to the playback side, while when plants were present in the arena, the moths switched preference to lay their eggs on the silent side. This latter choice was in accordance with their preference to lay eggs on thriving vs. dry plants while the first choice (without a plant) was somewhat surprising.

One explanation for this reversal in preference might be the multi-modal moth decision-making process. When drought-stressed signals alone (without a plant) were presented to the female moths, they might have become the only reliable signals for the presence of a plant in the arena, which can explain their strong preference for this side (Ramaswamy 1988; Sadek 2011; Zhang et al. 2024). In contrast, when we integrated thriving plants into the arena, the moths’ decision making became multi-factorial. Namely, on both sides of the arena, there were visual, texture, and olfactory cues of thriving plants, while the treatment side also exhibited an acoustic signal of a stressed plant. In this setup, the females’ oviposition preference was reversed to the side without the acoustic signal. This might suggest that the acoustic signal interpretation is content dependent, i.e., that the playback of stress sounds in a multi-factorial setup became a reliable signal of the physiological state of the plant. Therefore, the females reverted back to their original preference to oviposition on thriving plants.

To further examine this hypothesis, we conducted an additional experiment using the same protocol described for the “Sound gradient experiment” (see Methods), except that we placed a dehydrated plant (subjected to the stress treatment detailed in Experiment 1) on the side of the speaker that was playing plant-sounds. The resulting oviposition pattern closely mirrored those of our earlier studies: when presented with a stressed plant supposedly emitting dehydration sounds, *S. littoralis* females preferred to deposit their eggs on a dehydrated clicking plant rather than on a no-plant control (Supplementary Figure 4). These findings imply that a stressed, clicking plant is more attractive for oviposition than an empty substrate, suggesting that clicking might be a cue for the presence of a plant.

Supporting this hypothesis, in the two choice experiments the probability of laying eggs at all was significantly higher when a plant was present than in the absence of a plant. Specifically, eggs were laid on 68% vs. 54% of the nights with and without plants respectively (*p*=0.009; Binomial test comparing experiments two and three). The number of egg clusters was also higher when a plant was present (see Fig.1). We conclude that the moths were more reluctant to lay their eggs when no plant was present.

The preference for the silent plant vs. a plant with stress acoustic playback was not as clear as the preference for the thriving hydrated plants (compare Fig.1B1 and Fig.1B3). There are several potential explanations for this difference. First, moths probably rely on various cues, including olfaction, to detect a drying plant (Ramaswamy 1988; Sadek 2011; Zhang et al. 2024)). Although the playback allowed us to isolate the specific effect of the acoustic cue, and we tried to select equal plants, we could not control for other cues provided by the plant, and we may have provided the animal with a partial (and likely even contradictory) set of cues. For instance, the plants might have secreted drought-related volatiles and (although watered) might have occasionally emitted sounds spontaneously, reducing the effect of our playback. Indeed, a physiological measurement of plant volatiles suggested that drying plants can be (at least partially) distinguished by the moths (Fig.S3).

We further investigated the behavioral mechanism of the female moths as they explored the arena. We quantified the moths’ movement during the decision process in the experimental setup with drought-stressed acoustic signals played on one side, and with an equal-impedance resistor on the other side. Our findings indicated that their decision process typically included crossing over between the two sides of the arena and spending an increasing amount of time on the (drought-stressed) playback side. This suggests that females explore the available space and ultimately decide based on comparing the two.

Various plant species emit airborne ultrasonic clicks when they are drought-stressed, which can serve as reliable cues for the physiological condition of the plant (Khait et al. 2023). Our findings demonstrate that moths with auditory abilities use these clicks when choosing a site for oviposition. We hypothesize that some other species of insects might also exploit these acoustic cues to their advantage in different contexts. Pollinating insects, for example, might use drought-related sounds when choosing where to forage. Some insects might even be able to distinguish between clicks produced by different plants or under different conditions, such as drying plants vs. plants under a pathogen attack.

Plant clicks are ultrasonic and thus very different from most other outdoors sounds (such as wind sounds, as we also show in Khait et al. 2023). Moreover, because the clicks are ultrasonic and not very intense, they can only be picked up by the moths from a short distance (∼1.5m) which allows the moths to localize them in space.

The sounds emitted by drought-stressed plants are probably a cue rather than a signal, i.e., they did not evolve to convey information to insects. The interaction that we have demonstrated in this study therefore cannot be considered “communication” according to the conservative definition of the term, which relies on signals that have evolved to convey a specific message (Searcy and Nowicki 2005; Skyrms 2010). However, it is possible that some plants have evolved an ability to amplify their emissions or modify their spectral content to facilitate desirable interactions with animals and perhaps even with other plants (Veits et al. 2019). One exciting possibility would be that plants signal an insect attack by amplifying click intensity to recruit potential predators of the attacking insects, such as predatory insects, rodents, or bats. Such amplification could be achieved by various morphological modifications. Insects, on the other hand, might have evolved behavioral strategies to move near plants and pick up these weak acoustic signals. In conclusion, our study shows that moths are able to detect and respond to acoustic signals emitted by plants. This discovery suggests the existence of a third type of acoustic signal that moths utilize, in addition to those produced by bat echolocation and moth courtship clicks, raising new questions about the evolution of moth hearing. We predict that future studies will uncover more examples of acoustic communication between plants and animals.

## Methods

Experimental setup –We collected pupae of *Spodoptera littoralis* that were reared under controlled breeding conditions (reared on castor bean leaves, 25 ± 1 °C, 40% relative humidity with a 12–12 h light–dark cycle). Newly-emerged female and male moths were placed together until egg-laying was detected (approximately two days). Then we transferred the females to an experimental arena. Each arena was 100 x 50 x 50 cm3 in size, divided in the middle by a plastic partition half the height of the arena (Fig. 1A). On the partition, we placed a closed test tube with cotton wool containing 60% inverted sugar solution for ad libitum feeding throughout the experiment. Experiment 1 (see below) was performed in a greenhouse (2.5 x 4.5 x 3.5 m^3^) to simulate optimal conditions for plant development. The experiments involving acoustic signals (see below experiments 2,3,4,5 and 6) were performed in an acoustically shielded room (2.5×4×2.5 m^3^) to prevent acoustic interference. Each of the following treatments was performed simultaneously in up to four arenas. Moths could choose between the treatments presented on each side of the arena (see below) and oviposition was monitored daily for three days by counting the number of egg clusters. At the end of each night, we cleaned the arena of counted egg clusters using a cloth with ethanol, so that on the subsequent night, we would not expect there to be evidence of previous oviposition. We repeated the experiments under the same conditions until acquiring at least nine nights with egg-laying observations (eggs were not always laid, which is not surprising given the artificial conditions in the acoustic room used for these experiments). We refer to the cluster and not to the individual egg as the moth’s decision unit, because each cluster requires a decision about the location of oviposition, whereas the number of eggs could be affected by the general condition of the female or by external interference. Indeed, there was much variation in the number of eggs per cluster - 68 ±134 eggs (mean ± SE). However, to determine whether counting eggs would have altered our results, we conducted an experiment comparing cluster counts to individual egg counts (experiment 6). For experiments with actual plants, a young tomato plant (*Solanum lycopersicum*) in a small pot was used in all experiments. All the treatments are illustrated in Fig.1A-C. The number of repetitions of each treatment is noted in Table 1 and data is presented in supplementary Table 1. To maintain moths’ vitality through the experiment we have placed on the starting point (central platform) a closed test tube with cotton wool containing a 60% sugar solution for ad libitum feeding.

1. <u>Drought-stressed vs. well-hydrated plants</u>: We placed a single-stem tomato plant, 10 cm high, on either side of the arena. The plant on one side was drought-stressed (three days without watering), and the other was thriving and well-hydrated. Moths could lay eggs on either plant (Fig.1A).
2. <u>Playback of a drought-stressed plant vs. silence (without plants):</u> Each side contained an oviposition box (10 x 15 x 5 cm^3^ made of 0.5 x 0.5 cm^2^ mesh), covered with a paper towel. A speaker playing sounds recorded from a drought-stressed tomato plant (Khait et al. 2018) was placed under one of the two oviposition boxes (on one side of the arena). The speaker played drought sounds at the same intensity measured for real plants at a rate of 1 click per minute, with a stochastic 10% error in the intervals between clicks (see below for details on assessing intensity and playback rate). The oviposition box on the other side either had a resistor similar to the speaker in shape and identical in impedance to control for potential effects of the electric field created by the speaker (though we did not account for a magnetic field produced by the speaker, which might as well affect the choice), or no resistor (we did not find significant differences between the two silent controls, GLMM, p=0.58). The experiment was performed twice to strengthen the confidence in its results: the first trial was performed during August and September 2021 and the second during February to May 2022 (A pool of both trials and controls - with and without resistors - is presented in Fig.1B2). We also repeated this experiment a third time with a lower emission rate of 2 clicks per minute (Table 1).
3. Deaf females in a setup with silence vs. drought-stressed plant playback (without a plant): we deafened mated females by puncturing their tympanic membrane and placed them in an arena to assess their response to drought-stressed sounds, compared to a silent control (as described in experiment 2). Deafening surgical procedure: We performed a surgical procedure on female moths to deafen them. The procedure involved puncturing the tympanic membrane located at the thoraco-abdominal juncture using an entomological needle #2. The female moths recovered from the procedure within 2 minutes and were able to fly normally. We tested a sample of these females in a standard rearing box and found that they were able to lay eggs normally. To confirm that the surgery had successfully deafened the females, we conducted an inspection by playing a bat playback (the same as described below). We deafened a group of 20 moths and compared their reactions to a control group of 25 non-deafened moths. During the experiment, the moths were released in a dark acoustically isolated room (5.5 × 4.5 × 2.5 m3) with acoustic foam on the walls and ceiling and a single light source (12W mercury vapor bulb peaked at 1,650 lux), and while they were in flight around the light source, we emitted the sound. In the control group, 5 moths exhibited a response (such as falling or a significant change in direction), upon hearing the sound (scored by a naïve viewer who did not know whether the moths were treated). In contrast, none of the deafened moths displayed any reaction to the clicking stimulus (Q=4.5, p=0.03, Chi-square test).
4. Well-hydrated plants with and without playback of drought-stressed plant sound: There was an oviposition box on each side of the arena. One side played drought-stressed sounds while the other remained silent, with either a resistor or no sound (same as experiment number 2). Additionally, a thriving, healthy tomato plant was placed on each oviposition box. This experiment was performed twice, 12 months apart, to strengthen the confidence of its results (a pool of both trials and controls [with and without resistors] is presented in Fig.1B3). To determine the specificity of the response to plant sounds, two additional controls were performed:
5. Male moths: Five males were enclosed under the oviposition box with sugar water to maintain them. The control box had only sugar water without any moths (Fig.1C). We validated that males in this condition produced clicks by recording the sounds emitted by the five male moths enclosed overnight in an acoustically isolated container which showed that the males frequently click. The test was repeated five times and clicks were always emitted by the males (Supplementary Fig.1(.

### Playback

Drought sounds were recorded using an Hm16 Avisoft microphone and an HM116 Avisoft A/D from a distance of 10 cm in an isolated container with walls covered with acoustic foam (Khait et al. 2018). These recordings revealed emission intensities of at least 60 dB SPL (Re 20μPa) at a distance of 10 cm. The sounds were played using a Vifa speaker connected to an Avisoft D/A converter (Player 116).

We ensured that playback sound intensity was similar to that measured in real plants on the playback side of the arena (i.e., ∼60dB SPL at a distance of 10 cm) and that sound level on the control side was below the detection range of our system, that is, below 30dB SPL at 10 cm. We performed four calibration measurements using a calibrated GRAS 40DP microphone during the period of the experiments to validate that sound levels had not changed over time. Using the GRAS calibrated microphone, we also validated that the average sound intensity of the male moth sequences was the same as that of the playback plant sounds.

Validating the playback rate: The drying plant sounds in the box arenas (experiments 2-4, Fig.1B) were played back at a rate of 1 click per second (1 Hz) with up to 10% error in the intervals (caused by the computer controlling the system). This frequency is substantially higher than that found for a single young tomato plant (Khait *et al*. 2023). However, the rate that we played (60 clicks per minute) is ecologically relevant when considering a patch of tomato (or other) plants. To validate this, we aggregated 45 tomato seedlings in a planting tray (30 x 30 cm^2^) and placed the tray in an empty greenhouse. The plants were not watered for three days, and we recorded sound continuously for 50 hours (using the same Hm116 microphone setup noted above). When placing the microphone ∼20cm above the tray – as a flying moth would do, we measured a maximum click rate of 20 clicks per minute (i.e., 0.33 Hz). This is three-fold slower than the rate we used, but very similar to the rate that we used in the gradient experiment (see below). Moreover, when taking into account the moth’s detection range for this emission intensity which is likely ∼1.5 meters at least (Khait *et al*. 2023), a female moth could be exposed to a rate over three-fold higher (i.e., higher than 1 Hz) in a patch of drying plants (which would contain more than 100 seedlings in a typical bush of agricultural or wild hosts typical of this species). Notably, every plant that we examined was found to emit similar ultrasonic clicks when dehydrating (Khait *et al*. 2023), so this behavior could be relevant to other plants many of which grow as dense bushes.

### Sound gradient experiment

We used elongated arenas (150 × 20 × 5 cm^3^, Fig.2A). In the center of the arena (location 0), we placed a closed test tube with cotton wool containing a 60% sugar solution for ad libitum feeding (to maintain moths’ vitality). To facilitate accurate measurement of egg distances from the speaker (location −75), we printed a ruler and placed it along the bottom of the arena. Each moth was placed at the center of the arena at the beginning of the experiment. On the next morning, we recorded the locations of the egg clusters and counted the number of eggs in each cluster using a stereoscopic microscope, or a magnifying glass if the eggs were not laid on the ruler. Each female remained in the arena during the days starting three days after emerging from the pupa and mating, and until it died. After each night, we switched the locations of the speaker and the resistor within the arena. We measured a 30 dB SPL difference in intensity between the side of the speaker and the side of the resistor. The clicks were emitted at a frequency of 0.5 clicks per second (30 per minute).

### Tracking the females’ decision-making process

In order to investigate how moths survey the experimental arena and subsequently engage in a decision-making process, we conducted two additional trials in which we continuously recorded the movement of the moths throughout the night. In each trial, we placed four female moths on a platform in the middle of the arena, in which a speaker played drought-stressed plant sounds on one side, while on the other, control side we placed a silent resistor (as in treatment 3 above). We exchanged sides between trials and tracked the moths for six hours using an IR camera (Reolink RLC-511-5MP camera) placed above the arena. We then documented the position of each moth at 12 seconds intervals using the DLTdv 8 software (Hedrick 2008). Each individual was recognized according to its proximity to the last tracking point in order to reconstruct its full movement. We quantified how many times each individual crossed the center of the arena (the platform in the center was divided in the middle), and the proportion of time it spent in each side.

### Statistics

Mixed effect generalized linear models (GLMM) were used (in MATLAB) to examine the females’ choice of oviposition. Random effects were set as intercepts. The number of clusters was set as the explained variable. The treatment, i.e., playback or control, and the number of female moths in the arena, was set as a fixed effect. The number of the arena, the month in which the experiment was performed, the number of repetitions, and the night of the repetition were considered as random effects. Because we were analyzing counts (number of clusters), the model was run using a Poisson distribution. In the experiments in which we ran two repetitions of the same experiment, we added the session as another fixed parameter and we also ran the statistics separately for each session.

To deepen our understanding of the trends observed in the experiments, we implemented Bayesian model fittings for each choice-based experiment. In this analysis, “oviposition choices” were considered as distinct decisions. A value of 1 was assigned when the egg cluster was located on the side with the active speaker (or on the hydrated plant in the initial experiment) and a value of 0 was assigned for oviposition on the opposite side. We employed a Gaussian model, incorporating the number of females in each experiment as a random effect, with a prior mean of 0.5 and a standard deviation of 0.1. For each experiment, we sampled our data 16,000 times to calculate the posterior distribution from these samples. We used a Binomial GLMM to determine the effect of the treatment on the moths’ decision making. To achieve this, the proportion of time spent in each side of the arena was set as an explained variable, the playing side as a fixed effect, with the trial and the individual moth as random effects. To study the effect of time on the movement of the moths, we used Logistic GLMM in which the accumulated amount of time spent on the sound-playing side was set as an explained variable, the time as a fixed effect, and the trial and the individual moths were set as random effects.

To compare the distribution of eggs in the elongated arena to a random distribution, we generated an H0 distribution by randomly shuffling the locations of the speaker and resistor for each laid egg. This distribution was then compared to our actual egg count distribution using the Kolmogorov-Smirnov test (Supplementary Fig. 3).

## Acknowledgments

We thank Hadas Marcus for linguistic editing, Tal Erez for providing moths, Adi Segal, Einav Balachsan, Michael Martynenko, Shira Fraenkel, Yuval Lustig, and Morad Garam for experimental setup, Nitzan Shahar statistical consulting, Aya Goldshtein for reviewing the manuscript and statistical consulting, Mor Taub for graphic editing, Yonatan Nathan, and Inon Scharf for reviewing the manuscript.

## Supplementary

**Supplementary Fig 1.**
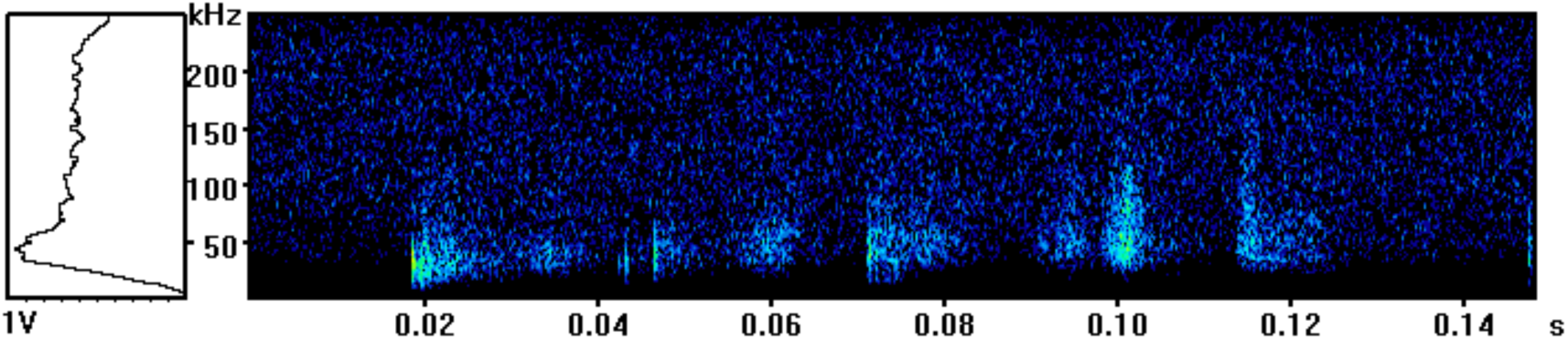
Male Egyptian cotton leaf moths *(S. littoralis)* courtship sequences recorded when we placed males in the arena (spectrogram presented).

**Supplementary Fig 2.**
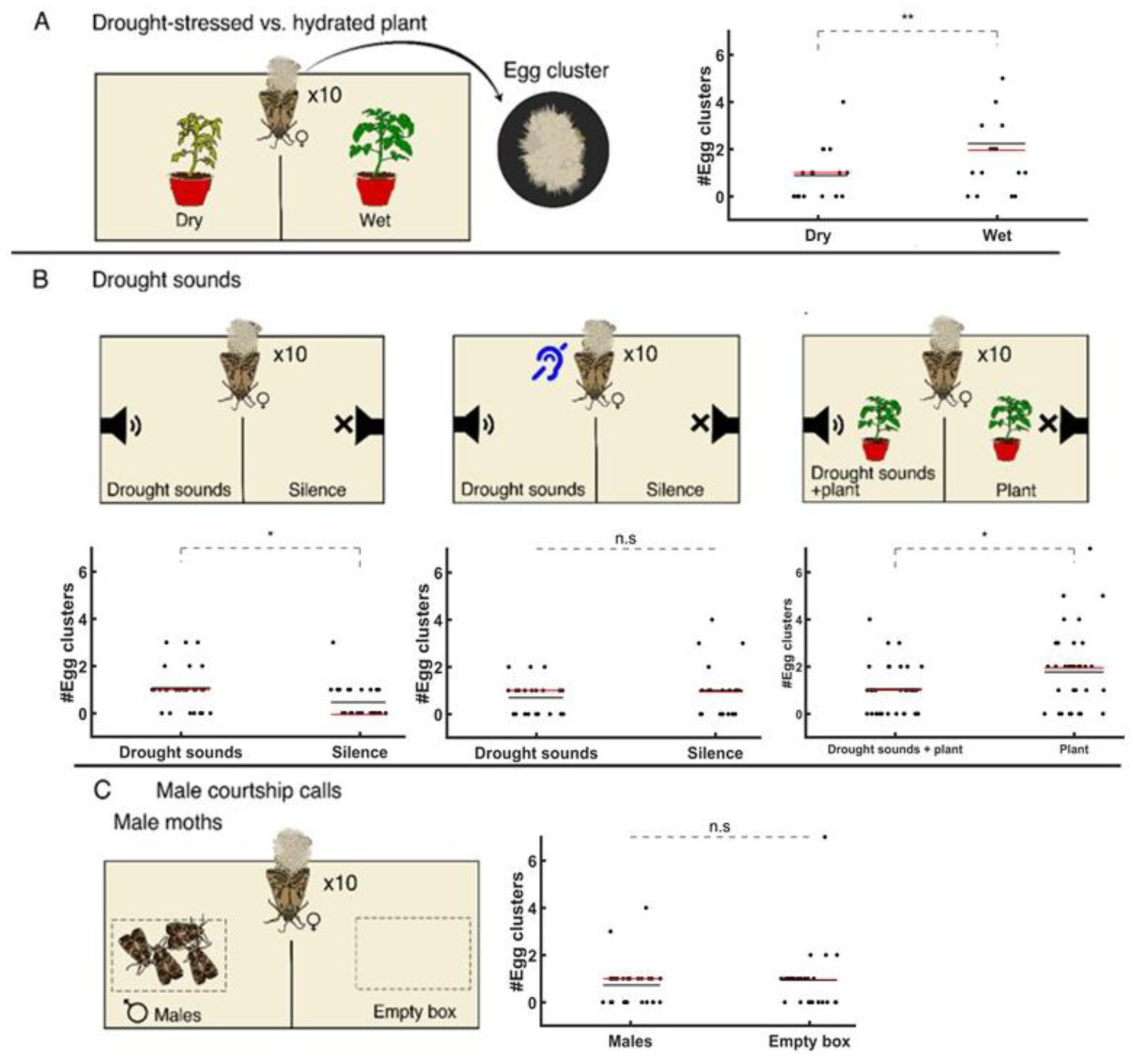
This figure replicates the experiment shown in Figure 1, displaying the raw measurements without Bayesian analysis. In all panels (A–D), the treatment is shown in the left section. These graphs summarize every datum collected throughout the two choices experiments. Each marker represents a cluster deposited at the choice indicated on the X-axis. The overall mean is overlaid as a solid black line, and the median as a solid red line. A) Drought-stressed vs. thriving plant (no playback). B1) Silence vs. drought-stressed plant playback (without a plant). B2) Deaf females in a setup with silence vs. drought-stressed plant playback (without a plant). B3) Silent plant vs. playback of drought-stressed plant. C) A box with male moths vs an empty box.

**Supplementary Fig 3.**
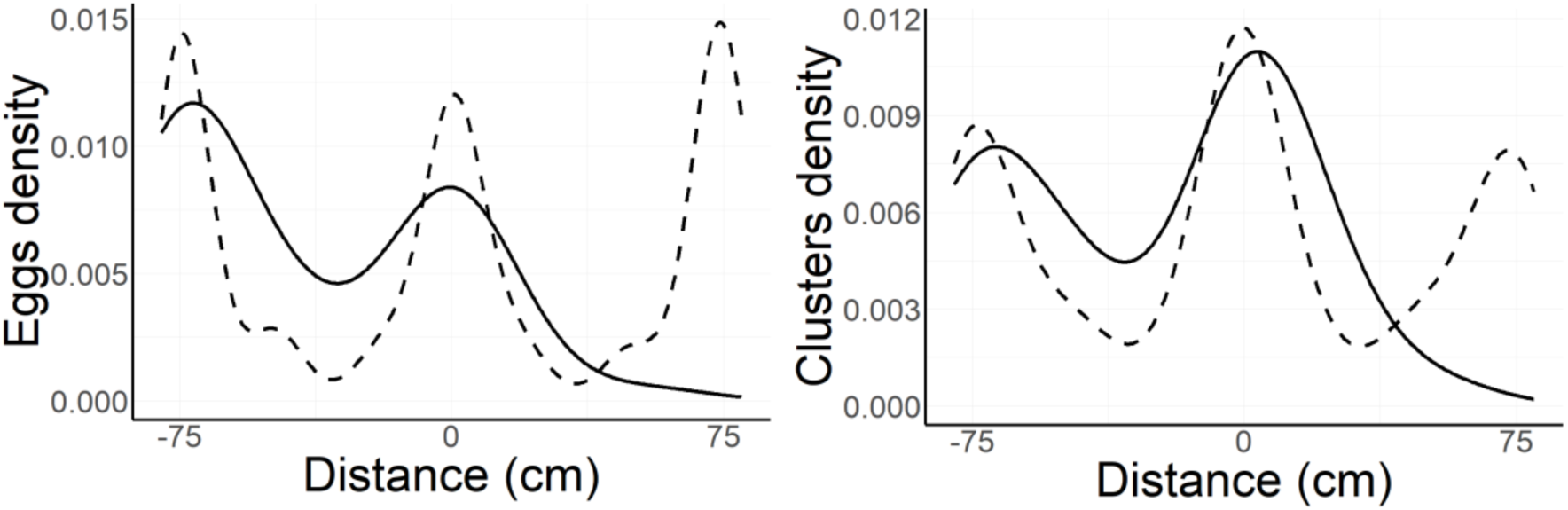
On the left, comparison between the egg count results (solid line) in the elongated arena and the pseudo-random distribution (dashed line) (K-S test, D = 0.3, p = 2.2 × 10⁻¹⁶). On The right, comparison between the clusters count results (solid line) pseudo-random distribution (dashed line) (K-S test, D = 0.21, p = 3.9 × 10⁻14). The speaker was placed on location −75, a feeder was placed on the center (location 0) and a resistor was placed on location 75. To exclude any potential effect of temporal correlations on egg laying, we have also rerun the statistics when only taking the first night, when the females laid clusters to avoid the desensitization or dependency. This test revealed similar results (D = 0.55, p-value < 2.2× 10⁻16).

**Supplementary Fig 4.**
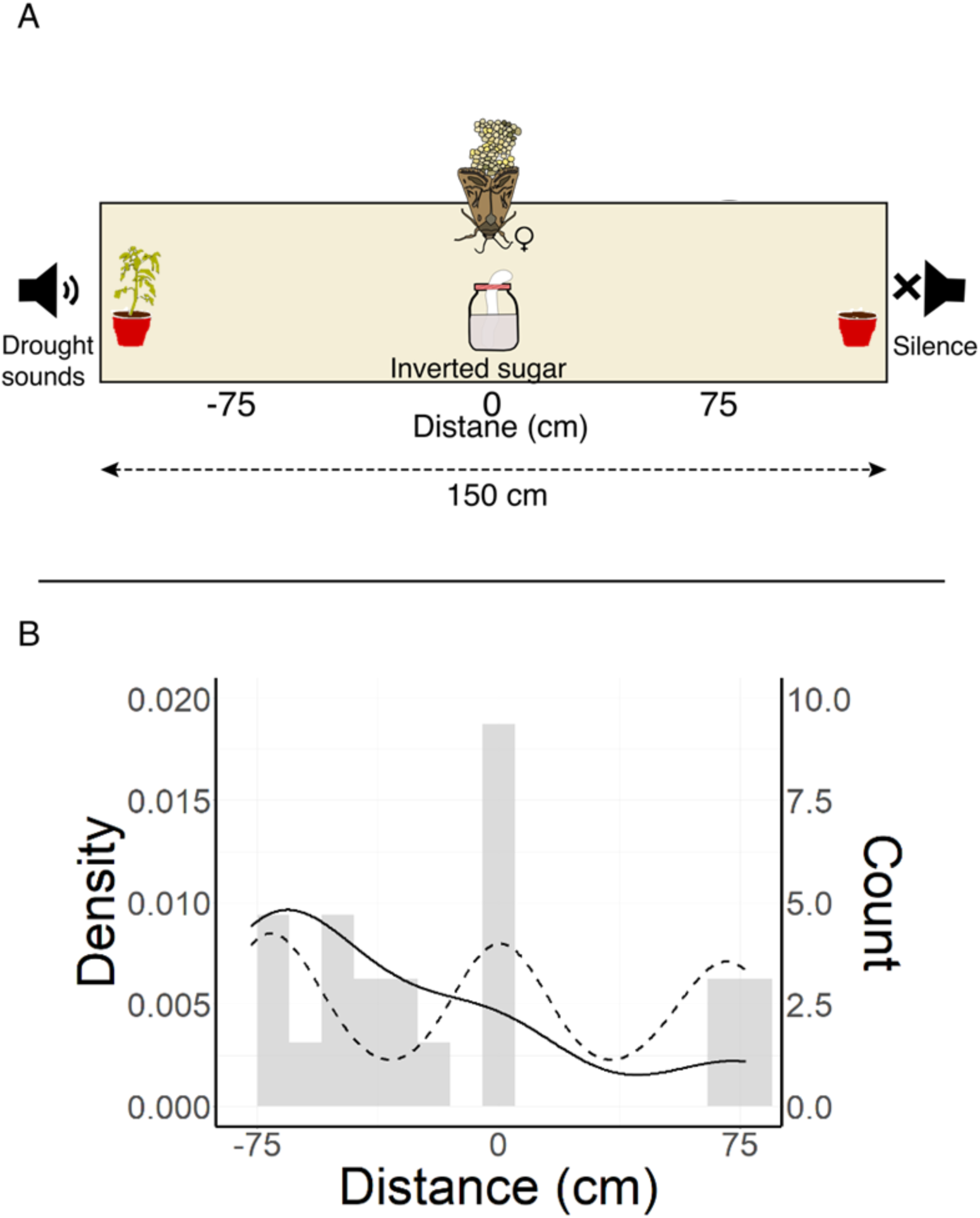
(A) We conducted an additional experiment using the same protocol described for the “sound-gradient experiment” (see Methods, Sound gradient experiment), except that we placed a dehydrated, plant (subjected to the stress treatment detailed in Experiment 1) on the speaker side, and a resistor plus soil on the control side. (B) The resulting oviposition pattern closely mirrored those of our earlier studies: when presented with a stressed plant versus an empty control, *S. littoralis* females deposited significantly more egg clusters on the dehydrated clicking plant. To test the effect of the treatments on the oviposition we compared the observed cluster locations (solid line) to pseudo-random distribution (dashed line). We found significant differences between the two distributions (K-S test, D = 0.29, p = 0.001). The speaker was placed at location −75cm, a feeder was placed on the center (location 0) and a resistor was placed at location 75cm. We have also rerun the statistics when only taking the first night when the females laid clusters to avoid the fear of dependency. These tests revealed similar results (D = 0.34, p = 0.020). Light-gray bars denote the observed measurements aggregated into 10 cm bins (N=20).

